# Limb-clasping, cognitive deficit and increased vulnerability to kainic acid - induced seizures in neuronal GPI anchor deficiency mouse models

**DOI:** 10.1101/2020.10.21.348334

**Authors:** Lenin C. Kandasamy, Mina Tsukamoto, Vitaliy Banov, Sambuu Tsetsegee, Yutaro Nagasawa, Mitsuhiro Kato, Naomichi Matsumoto, Junji Takeda, Shigeyoshi Itohara, Sonoko Ogawa, Larry J. Young, Qi Zhang

**Affiliations:** Laboratory of Social Neural Networks, Center for Social Neural Networks, University of Tsukuba, Japan; Laboratory for Behavioral Genetics, CBS, RIKEN, Japan; Institute of Neuroinformatics, University of Zürich, ETH Zürich, Switzerland; Department of Pediatrics, Showa University School of Medicine, Japan; Department of Human Genetics, Graduate School of Medicine, Yokohama City University, Japan; Yabumoto Department of Intractable Disease Research, Research Institute for Microbial Diseases, Osaka University, Japan; Laboratory of Behavioral Neuroendocrinology, Faculty of Human Sciences, University of Tsukuba, Japan; Faculty of Human Sciences, Center for Social Neural Networks, University of Tsukuba, Japan; Center for Translational Social Neuroscience, Department of Psychiatry and Behavioral Sciences, Yerkes National Primate Research Center, Emory University, Atlanta GA 30329

**Keywords:** GPI, PIGA, conditional knockout, excitatory, inhibitory, memory, seizure

## Abstract

Post-translational modification of a protein with glycosylphosphatidylinositol (GPI) is a conserved mechanism exists in all eukaryotes. Thus far, more than 150 human GPI anchored proteins have been discovered and about 30 enzymes have been reported to be involved in the biosynthesis and maturation of mammalian GPI. Phosphatidylinositol glycan biosynthesis class A protein (PIGA) catalyzes the very first step of GPI anchor biosynthesis. Patients carrying a mutation of the *PIGA* gene usually suffer from intractable epilepsy and intellectual developmental disorder. We generated three mouse models with PIGA deficits specifically in telencephalon excitatory neurons (Ex-M-cko), inhibitory neurons (In-M-cko), or thalamic neurons (Th-H-cko), respectively. Both Ex-M-cko and In-M-cko mice showed impaired long-term fear memory and were more susceptible to kainic acid (KA)-induced seizures. In addition, In-M-cko demonstrated a severe limb-clasping phenotype. Hippocampal synapse changes were observed in Ex-M-cko mice. Our *Piga* conditional knockout mouse models provide powerful tools to understand the cell-type specific mechanisms underlying inherited GPI deficiency and to test different therapeutic modalities.

## Introduction

The international focus on human congenital disorders of “sugar coding”-glycosylation (CDG) has grown considerably in the past 10 years along with great advances in next generation sequencing technology. A new glycosylation disorder was reported, on average, every 17 days in 2013 (1). One important classification of CDG is glycosylphosphatidylinositol (GPI) anchor biosynthesis defects (2). GPI anchoring of proteins (GPI-AP) is a conserved post-translational modification among all eukaryotes, and at least 150 human proteins are GPI-APs, including receptors, adhesion molecules, complement regulators and enzymes (3). Biosynthesis and maturation of mammalian GPI-APs requires about 30 genes (3). Germline mutations of any of these genes could cause inherited GPI deficiency (IGD) with a wide spectrum of symptoms, including pronounced neurologic impairments, such as intellectual disability and intractable seizures, early onset epileptic encephalopathies (EOEE), cerebral and/or cerebellar atrophy, hypotonia, motor incoordination and ataxia, cortical visual impairment and sensorineural deafness (4).

Phosphatidylinositol glycan biosynthesis class A protein (PIGA) catalyzes the very first step of GPI anchor biosynthesis and the loss function of the gene encoding PIGA (*PIGA*) completely abolishes GPI anchor production (5). The group of Matsumoto and others have applied whole-exome sequencing and identified multiple *PIGA* germline mutations in patients with neurological symptoms from 15 unrelated families (6–15). Therefore, it would be important to study the detailed molecular, cellular and neural circuit mechanisms underlying the IGD encephalopathy by using *Piga* mutated animal models. Complete disruption of the *Piga* gene caused early embryonic lethality (16). Takeda and Kinoshita developed a conditional *Piga* ^flox^ mouse line, in which exon 6 of *Piga* is flanked by two identically orientated loxP sites, and proved that *Piga* function is essential for proper skin differentiation and maintenance (17). More recently, this conditional knock out (cko) system was used in *Nestin-Cre* mice to delete *Piga* in the neuroepithelial stem cell population that differentiates into neurons, astrocytes and oligodendrocytes. These mutated mice showed deficits in cerebellar and white matter development, ataxia and tremor, and did not survive past weaning. The early lethality of these mice exclude the possibility to examine their cognitive and epileptic phenotype thoroughly (18). In addition, *Nestin* is also expressed in a variety of tissues and progenitor cells, including pancreatic islets, skeletal muscle satellite cells, developing myotomes, testis, hair follicle, heart, and the non-hematopoietic fraction of the bone marrow (19), further adding to the complexity of explaining the mutants’ phenotype.

In the current study, by using three Cre recombinase (*Cre*) mouse lines including *Emx1-Cre*, *Vgat-Cre*, and *Pkcd-Cre* lines, we successfully ablated *Piga* in three populations of neurons: telencephalon excitatory neurons; inhibitory neurons and thalamic neurons, respectively. As *Piga* is localized on the X chromosome, a male mouse bearing a floxed *Piga* (*Piga*^flox/Y^) has only targeted *Piga* locus, and will completely disrupt *Piga* in *Cre*-expressing cell lineages (hemizygous cko). Whereas a female mouse bearing a loxP site (*Piga*^flox/+^) will have *Piga* partially ablated in *Cre*-expressing cell lineages due to random X chromosome inactivation (mosaic cko). *Emx1-Cre* hemizygous cko mice (Ex-H-cko) and *Vgat-Cre* hemizygous cko mice (In-H-cko) failed to survive after birth. *Emx1-Cre* mosaic cko mice (Ex-M-cko), *Vgat-Cre* mosaic cko mice (In-M-cko) and *Pkcd-Cre* hemizygous cko mice (Th-H-cko) survived to adulthood. Further analysis revealed that Ex-M-cko and In-M-cko mutants performed poorly in fear memory task, and their susceptibility to kainic acid (KA) induced seizures was significantly increased. In addition, In-M-cko mutants demonstrated a robust limb clasping phenotype and decreased bodyweight. Our animal models with GPI deficit in different populations of neurons provided a useful tool to screen effective treatments of IGD patients.

## Results

### Generation of different neuronal population-specific *Piga* cko mice

In the *Piga* ^flox^ mouse line, three loxP sites with identical orientation were inserted within the *Piga* gene on X chromosome, with one loxP localized in intron 5 and the other two loxP sites flanking a *Neo* gene localized downstream of exon 6 (Fig1.A). Previous work using this mouse line proved that deletion of exon 6 upon *Cre* recombination resulted in an efficient ablation of *Piga* and disrupted the cell surface expression of GPI-APs (16–18, 20). Our *In situ* hybridization on adult mouse brain showed that *Piga* is widely expressed in almost all the neurons, and has a very strong expression in CA1 pyramidal cells and dentate gyrus (DG) granule cells of the hippocampus, and piriform cortex (Fig1.B). To investigate the functions of different neuronal populations in the encephalopathy of IGD, we crossed *Piga* ^flox/+^ female mice with the following three *Cre* lines of male mice: Emx1-Cre line, which induces recombination in excitatory neurons in the cerebral cortex and limbic structures (Fig1.C) (21); *Vgat-Cre* line, which induces recombination in all inhibitory neurons throughout the brain (Fig1.D) (22); and *Pkcd-Cre* line, which induces recombination in almost the entire thalamus (Fig1.E) (23). As *Piga* is localized on the X chromosome, theoretically by this breeding strategy we could get male hemizygous cko mice with the genotype of “*Piga* ^flox/Y^, Cre+”, and female mosaic cko mouse with the genotype of “*Piga* ^flox/+^, Cre+”. The Emx1-Cre line is a knock-in mouse, we used the homozygous Emx1^Cre/Cre^ so that all the offspring carried the *Cre* gene. The *Vgat-Cre* line and *Pkcd-cre* line are BAC transgenic lines. The *Vgat-Cre* mice we used harbor homozygous transgenic alleles so that all the offspring expressed Cre. The *Pkcd-cre* mice we obtained harbor heterozygous transgenic alleles thus the offspring could be either positive or negative for *Cre*. We performed PCR analysis of tail DNA from 22-day-old offspring to genotype both the *Piga* alleles and *Cre*. We obtained double positive female mice, *Piga* ^flox/+^,*Cre+*, from the breeding pairs of all the three *Cre* lines, meaning that mice with mosaic conditional knockout of *Piga* in excitatory neurons (Ex-M-cko), inhibitory neurons (In-M-cko), and thalamus neurons (Th-M-cko) could survive after weaning. We never obtained *Piga* ^flox/Y^,Cre+ male mouse from the *Emx1-Cre* and *Vgat-Cre* breeding groups, meaning that complete deletion of *Piga* in excitatory neurons or inhibitory neurons led to early death during embryo stage or soon after birth. However, we did get *Piga* ^flox/Y^,Cre+ male mice from *Pkcd-cre* breeders, meaning that the mutants with *Piga* totally disrupted in thalamus (Th-H-cko) could survive after weaning. Unless specified, in all the experiments, Ex-M-cko and IN-M-cko mutants were compared with their female littermates (*Piga*^+/+^,Cre+), Th-H-cko mutants were compared with their male littermates (*Piga*^flox/Y^,Cre-). Table 1 shows the genotyping results from the first-round crossing of all the three breeding groups. We further examined E13.5 embryos from two *Piga* ^flox/+^ X *Emx1-Cre* breeding pairs and three *Piga* ^flox/+^ X *Vgat-Cre* breeding pairs. From 22 embryos of *Emx1-Cre* breeders, there were 2 embryos that showed obvious microcephaly and their genotype was *Piga* ^flox^,Cre+ (Fig1.F and G). From 32 embryos of the *Vgat-Cre* breeders, 3 embryos displayed an amorphous sphere morphology with the genotype *Piga* ^flox^,Cre+ (Fig1. H and I).

**Figure1.**
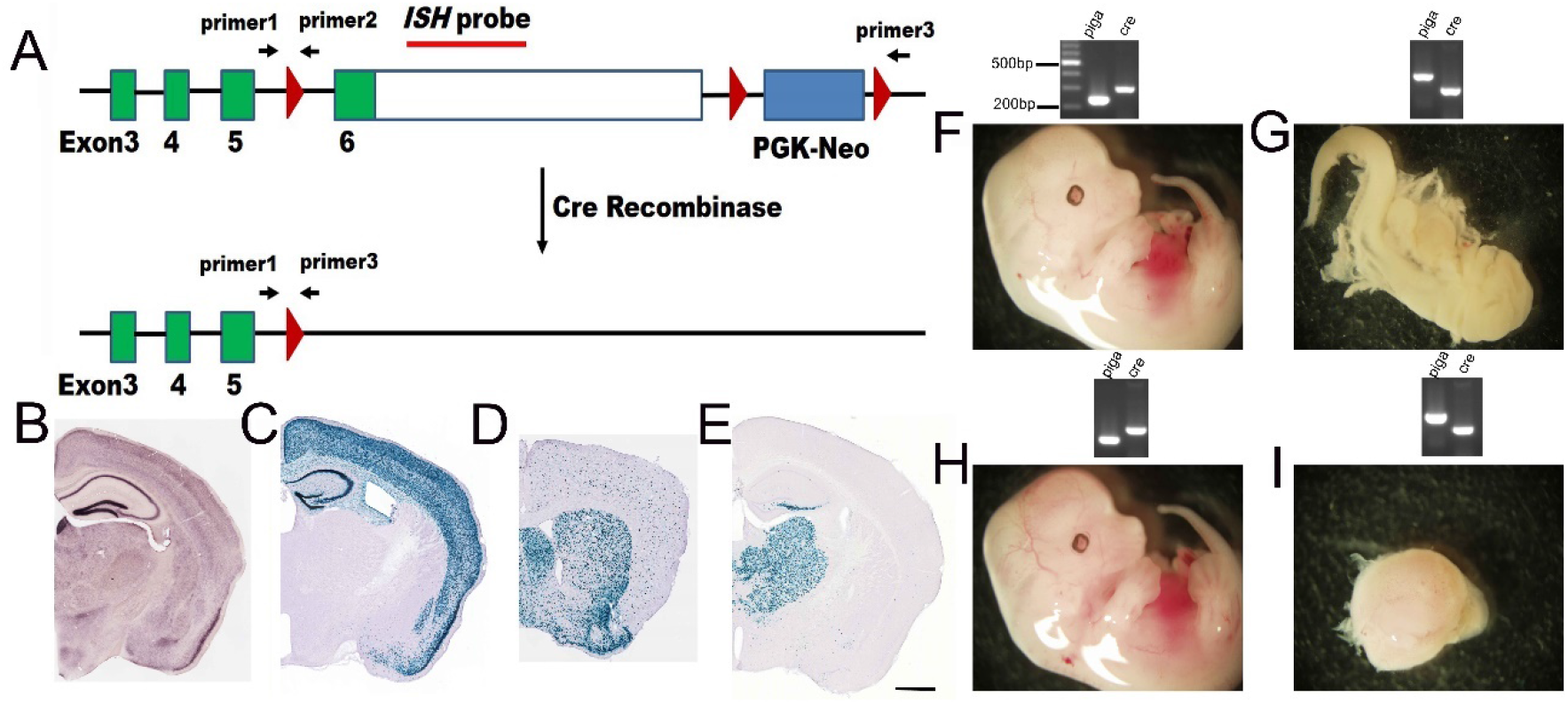
Creation of *Piga* conditional knockout mice. (A) The construction of *Piga*^flox^ mouse. The binding sites of three primers for genotyping and detection of *Piga* disruption, and that of *in situ* hybridization probe are indicated. (B) I*n situ* hybridization of *Piga* in adult mouse brain. (C-E) The X-gal staining of Rosa-NLS-LacZ reporter mice crossed with Emx1-Cre (C), *Vgat-Cre* (D), and Pkcd-Cre (E) lines, respectively. (F-I) E13.5 embryos with the genotype of *Piga*^WT^,*Cre+*(F); *Piga*^flox^, Cre+ (G) from the Emx-Cre breeding group. And E13.5 embryos with the genotype of *Piga*^WT^,Cre+(H); *Piga*^flox^, Cre+ (I) from the *Vagt-Cre* breeding group. The PCR result of genotyping from tail DNA (for I, the DNA was extracted from a piece of the out layer of the armorphous sphere due to deficiency of a tail) are shown above each image of the embryo. Scale bar:1mm

**Table1.**
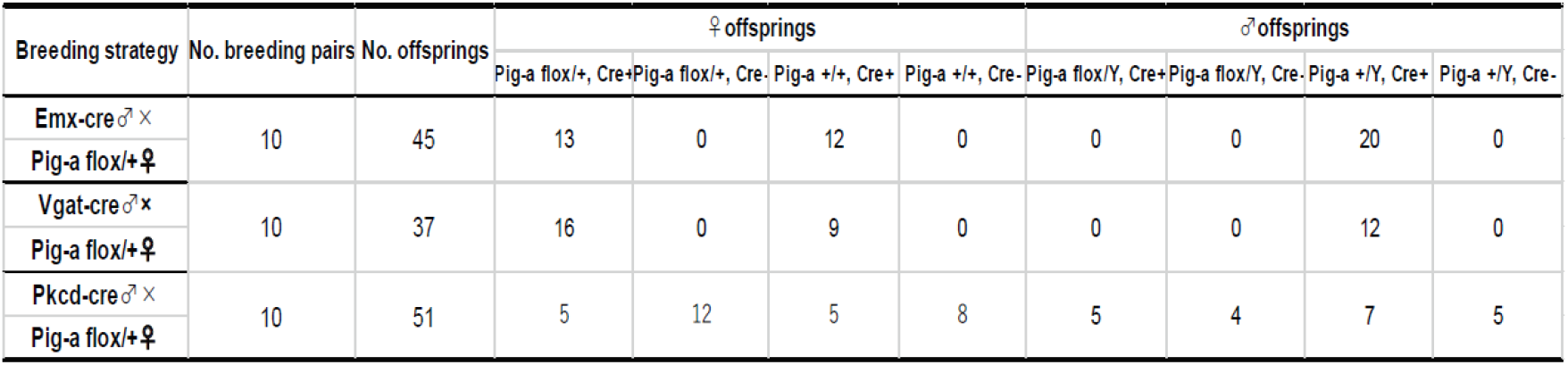
The mosaic and hemizygous mutants obtained from each cre lines in the 1^st^ round breeding.

We are interested in possible phenotypes resembling intellectual disability and epilepsy, therefore we focused on the cko mice that can survive after weaning. To confirm the deletion of *Piga* in specific targeted neuronal populations, we obtained neurons from hippocampus, striatum, thalamus and cortex using a well-established NeuN-labeled FACS method (24) and performed PCR using three primers shown in Fig1.A. The bands for “wild-type”, “targeted”, and “disrupted” *Piga* are about 250bp, 420bp and 550bp, respectively. Our results showed that in Ex-M-cko mutants, where *Cre* targeted *Piga* ^flox^ in excitatory neurons, the “disrupted” band from hippocampus was the most prominent, which is in consistent with the fact that the hippocampus GABAergic inhibitory interneurons represent only ~10–15% of the total neuronal population (25) (Fig2.A). In contrast, in In-M-cko where *Cre* targeted *Piga* ^flox^ inhibitory neurons, the “disrupted band” from striatum was the most prominent, consistent with the fact that 95% of neurons in striatum are GABAergic inhibitory neurons (26) (Fig2.A). In Th-H-cko, as expected, there was only a disrupted band from thalamic neurons but not from those from other brain regions (Fig2.B). In addition, there was no “disrupted” band from tails of the mutants, indicating the specificity of our cko system. To further confirm the specificity and efficiency of *Piga* ablation, we generated a probe that covers about 790bp of non-coding region of *Piga* exon 6. *In situ* hybridization on Th-H-cko tissue showed a dramatic decrease of signal in thalamus compared to that from a control littermate, whereas the signal in hippocampus and cortex was comparable with that from the control (Fig2.C,D). NetrinG1 is synaptic adhesion GPI-AP and it is mainly distributed at synaptic sites in adults (27, 28). We did immunohistochemistry for netrinG1 on Th-H-cko mutants. The signal of netrinG1 on cortex layer 4 was completely lost, whereas the signal in hippocampus CA1 stratum lacunosum moleculare and DG outer molecular layer was comparable with that from the control (Fig2.E,F). This result demonstrated that GPI synthesis deficit in thalamus neurons disrupted the synaptic delivery of netrin-G1 through thalamocortical axons, while the delivery of netrinG1 from entorhinal cortex to hippocampus through temporoammonic axons and lateral perforant axons was intact. In summary, by PCR of the disrupted *Piga* product, *in situ* hybridization of *Piga*, and examination of the surface delivery of a known GPI-AP, we provide convincing evidence that the ablation of *Piga* was specific in different neuronal populations.

**Figure2.**
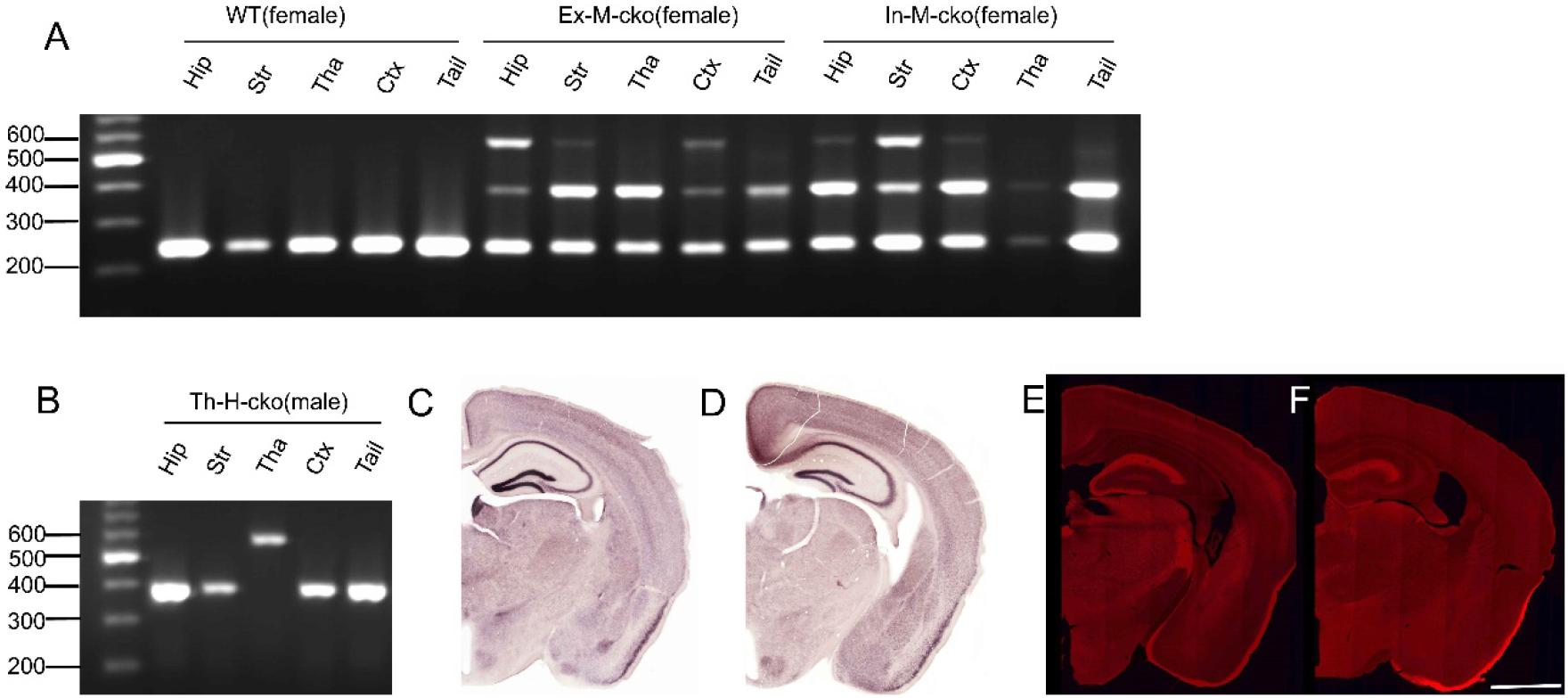
Specific deletion of *Piga* in different neuronal populations. (A, B) PCR was performed using neuronal DNA from different brain regions with all the three primers (Primer1 to Primer3) indicated in Fig1A.(C,D) *In situ* hybridization of *Piga* with Th-H-cko(C) and *Piga*^flox^, Cre-male control(D).(E,F) Immunohistochemistry of netrin-G1 with Th-H-cko(E) and *Piga*^flox^, Cre-male control (F). Scale bar: 1.8mm

### The deficit of PIGA in inhibitory neurons resulted in weight loss and limb clasping

We examined the phenotypes of Ex-M-cko, In-M-cko, and Th-H-cko mutants. Ex-M-cko, In-M-cko mutants were compared with *Piga* ^+/+^,Cre+ female control littermates. Th-H-cko mutants were compared with “*Piga* ^flox/Y^, Cre-” male control littermates. All these mutants could survive as long as their control littermates, at least for one year. The most distinctive difference was that all In-M-cko mutants showed smaller body size (Fig3.A). We compared the body weight of 7-month old mice (n=12/group), a one-way ANOVA showed there was a significant difference among female Ex-M-cko, In-M-cko and *Piga*^+/+^,Cre+ control (Fig3.F), F(2,33)=59.35, P<0.0001. Post hoc comparisons using the Bonferroni test indicated that the mean value for In-M-cko (M=18.28g, SE=0.64) was significantly lower and about 60% of that of the controls (M=28.82g, SE=0.71, P<0.0001). The mean body weight for Ex-M-cko (M=25.06g, SE=0.72) was slightly but statistically significantly lower than controls as well, P<0.01. An unpaired t-test was performed to compare the weight between male Th-H-cko mutants and “*Piga* ^flox/Y^, Cre-” controls (Fig3.G). There was no significant difference in the weight of Th-H-cko (M=34.73, SE=0.75) and control mice (M=33.85, SE=0.67; t(22)=0.099, p=0.756).

**Figure3.**
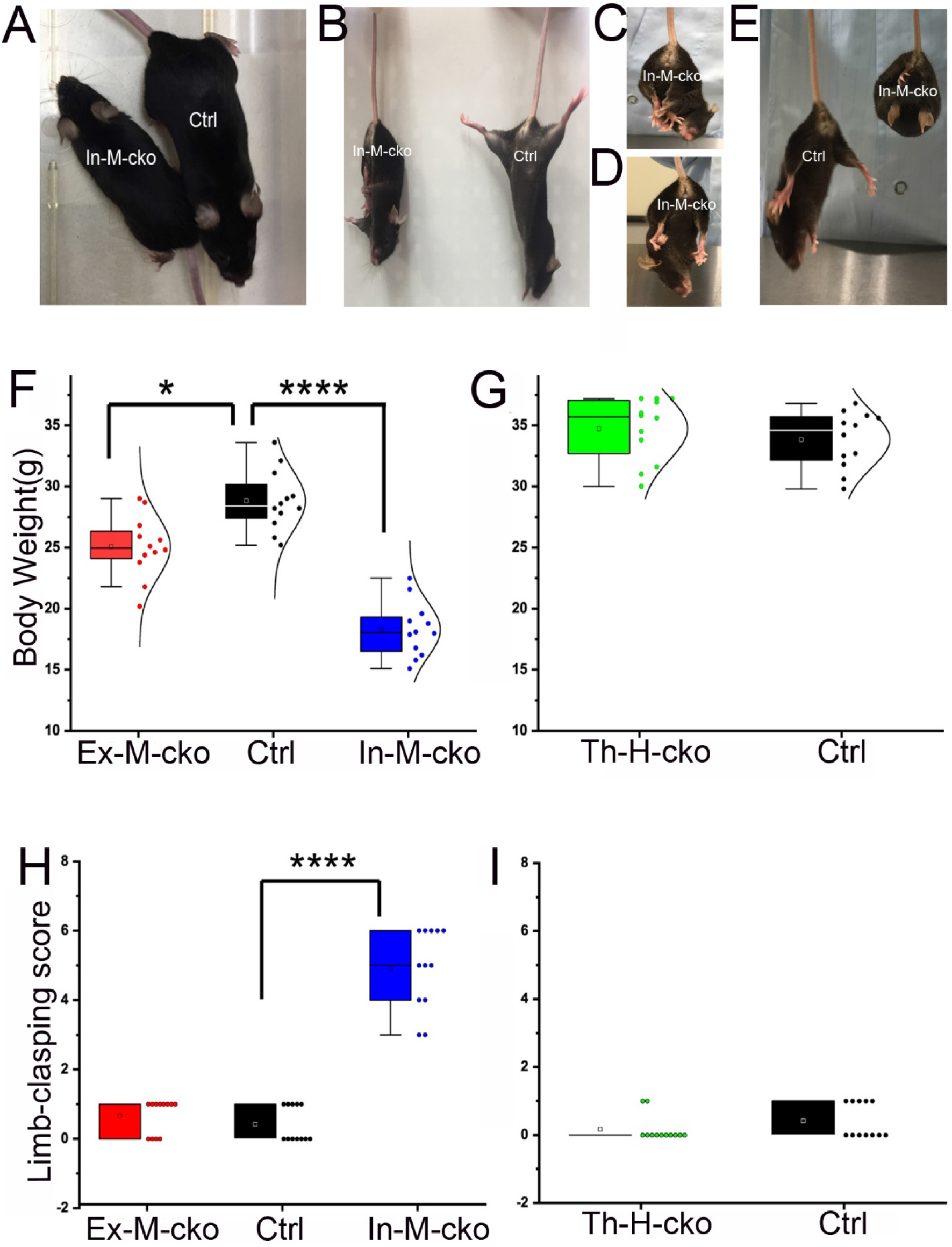
The body weight and limb-clasping phenotype in *Piga* mutants. (A-C) Body size (A) and weight (B,C) was compared between *Piga* mutants and their same sex controls. (D-I) Limb-clasping phenotype in In-M-cko mutants. In-M-cko mutants clasped three limbs (D), four limbs (E) or fore-hind limbs (F) together, and showed a bat-like posture (G). The limb clasping score was compared between *Piga* mutants and their same gender controls (H,I). In the box charts, the boxes are determined by the 25th and 75th percentiles. The whiskers are determined by the 5th and 95th percentiles. Mean and medium values are shown as the small square and line inside the boxes. Each individual value is plotted beside the box (B,C,H,I).

Another prominent phenotype of In-M-cko mutants was limb clasping (Fig3.B-D) and a bat-like posture (Fig3.E) when picked up by their tails. In fact, all the In-M-cko mice demonstrated obvious limb clasping. We evaluated the limb clasping of each cko group (n=12/group). A Kruskal-Wallis H test showed that there was a statistically significant difference in limb clasping score among female Ex-M-cko, In-M-cko and *Piga*^+/+^,Cre+ controls (Fig3.H),(χ2(2) = 29.40, P<0.0001), with a mean rank limb clasping score of 16 for Ex-M-cko, 30.5 for In-M-cko and 9 for controls. A post hoc test adjusted by Bonferroni correction showed that there was a significance between In-M-cko and control, P<0.0001, whereas the difference between Ex-M-cko and control did not reach statistical significance, P=0.25. A Mann-Whitney U test was applied to compare the limb clasping between male Th-H-cko mutants and “*Piga* ^flox/Y^, Cre-”control (Fig3.I). There was no significant difference between the mean rank of Th-H-cko (M=11) and that of control (M=14; U=54, p=0.19). All of the mutants including In-M-cko demonstrated normal gaits, and none of these mutants showed ataxia and tremor, at least by the age of 1 year old. In summary, mosaic ckos from either excitatory or inhibitory neuronal populations showed reduced bodyweight, and this weight reduction was especially remarkable in In-M-cko mutant. In addition, In-M-cko mutant demonstrated a robust limb-clasping phenotype.

### The deficit of PIGA in either inhibitory neurons or excitatory neurons impaired fear memory and caused an enhanced vulnerability to kainic acid (KA)-induced seizures

Prominent clinical features of IGD patients are intellectual disability and epilepsy. We examined whether our cko mice developed similar phenotypes. We used a classical contextual and cued fear conditioning task to test learning and memory ability of mutants (n=11/group). On Day1, a mouse learns to associate a context and an auditory cue to electric shocks. The next day they are tested to see if they remember the context associated with aversive stimuli, and the 3^rd^ day they are tested to see if they remember the tone paired with a shock (Fig4.A). A Mixed Factorial ANOVA was performed to compare the freezing level on each day of Ex-M-cko, In-M-cko and *Piga*^+/+^,Cre+ controls (Fig4.B). For Day1, conditioning day, Macuchly’s test indicated that the assumption of sphericity had been violated so that Greenhouse-Geisser correction was used, there was no significant interaction between genotype and time: F(5.25, 78.80)=0.70, P=0.63; and there was no significant difference for genotype: F(2, 30)=1.02, P=0.374. For Day2 contextual memory test day, Mauchly's test indicated that the assumption of sphericity had not been violated, χ2(9) = 7.9, P=0.54, there was no significant interaction between genotype and time: F(8,120)=0.965, P=0.47; however, the main effect for genotype was significant: F(2, 30)=13.8, P<0.0001. Bonferroni corrected post hoc test showed that the freezing level of In-M-cko mutants was significantly lower than that of control, P<0.0001; and the freezing level of Ex-M-cko showed a tendency to be lower than that of control but did not reach statistical significance, P=0.058. The multiple comparison of mean values (corrected by Bonferroni test) at each time point among three genotypes is shown in Table 2A and B. For Day3 cued memory test day, there was no significant difference among three groups: F(2, 30)=0.30, P=0.74. A Mixed Factorial ANOVA was performed to compare the freezing level on each day of male Th-H-cko mutants and “*Piga* ^flox/Y^, Cre-” (Fig4.C). For Day1, there was no significant difference between the two groups, F(1,18)=1.69, P=0.21. For Day2, there was no significant difference between the two groups, F(1,18)=0.49, P=0.49. For Day3, F(1,18)=0.01, P=0.92. The pain sensation of all the three mutants were comparable with their control mice (data not shown). Our results revealed that deficit of PIGA in either inhibitory or excitatory neurons decreased the long-term memory of contextual fear, consistent with the important role of GPI-APs in cognitive functions.

**Figure4.**
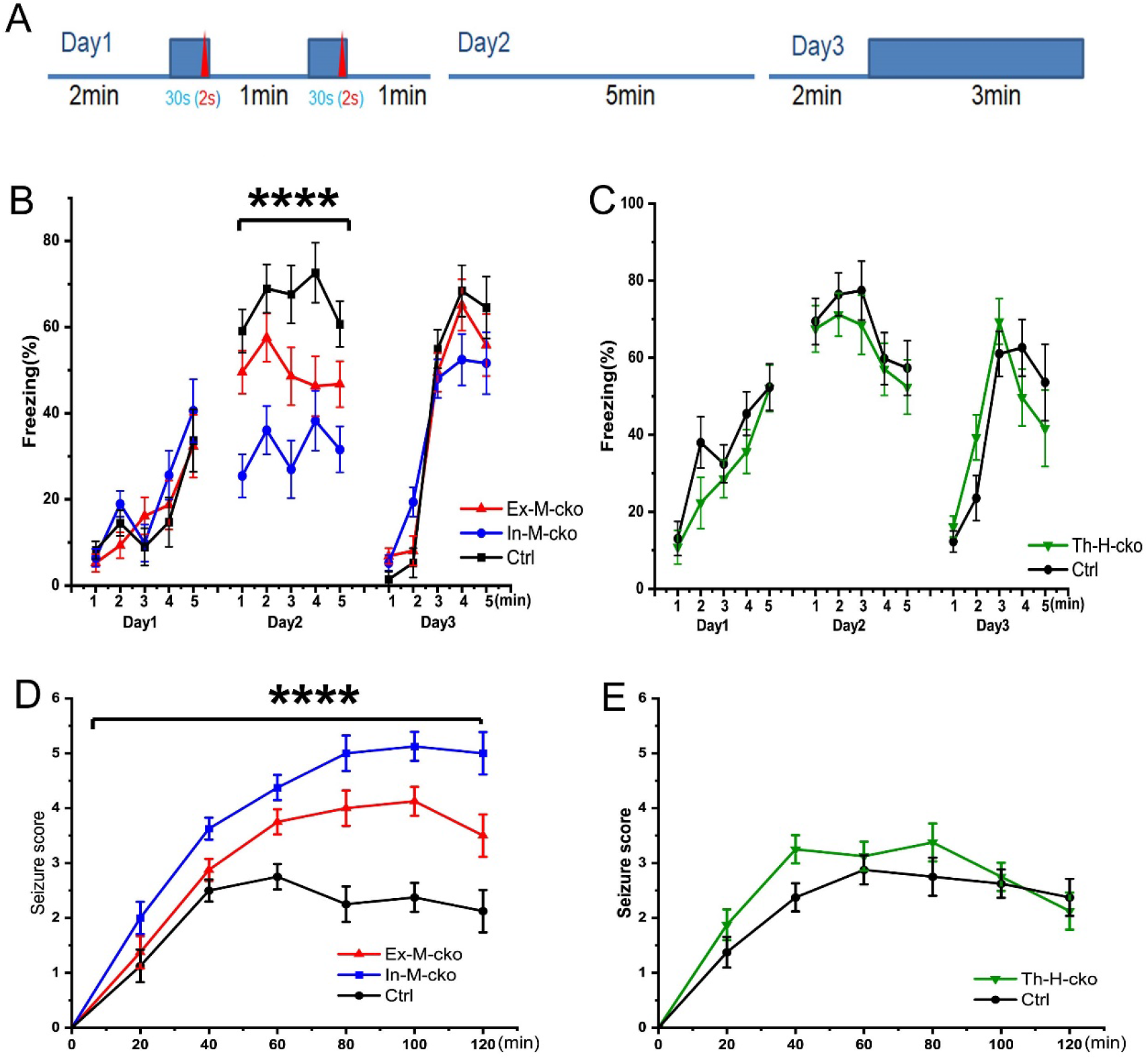
*Piga* mutants showed altered cognition and susceptibility to KA induced seizures. (A) A diagram of fear conditioning test. (B,C) The freezing percentage was compared between *Piga* mutants and their same sex controls. (D,E) The behavioral seizure score was compared between *Piga* mutants and their same sex controls. In the line charts, Mean ± SE are indicated for each time point.

**Table2.** The simple effects test result of the contextual fear memory among Ex-M-cko, In-M-cko and control. (A)The Bonferroni test corrected p value of each comparison between the freezing percentage of two genotype groups at each time point in contextual fear memory task. (B) The mean freezing percentage value of each genotype at each time point in contextual fear memory task.

| time(min) |  |  | Sig.<sup>b</sup> |
| --- | --- | --- | --- |
| 1 | Ex-M-cko | In-M-cko | .006 |
|  |  | Ctrl | .569 |
|  | In-M-cko | Ex-M-cko | .006 |
|  |  | Ctrl | .000 |
|  | Ctrl | Ex-M-cko | .569 |
|  |  | In-M-cko | .000 |
| 2 | Ex-M-cko | In-M-cko | .033 |
|  |  | Ctrl | .486 |
|  | In-M-cko | Ex-M-cko | .033 |
|  |  | Ctrl | .001 |
|  | Ctrl | Ex-M-cko | .486 |
|  |  | In-M-cko | .001 |
| 3 | Ex-M-cko | In-M-cko | .091 |
|  |  | Ctrl | .164 |
|  | In-M-cko | Ex-M-cko | .091 |
|  |  | Ctrl | .001 |
|  | Ctrl | Ex-M-cko | .164 |
|  |  | In-M-cko | .001 |
| 4 | Ex-M-cko | In-M-cko | 1.000 |
|  |  | Ctrl | .036 |
|  | In-M-cko | Ex-M-cko | 1.000 |
|  |  | Ctrl | .004 |
|  | Ctrl | Ex-M-cko | .036 |
|  |  | In-M-cko | .004 |
| 5 | Ex-M-cko | In-M-cko | .159 |
|  |  | Ctrl | .221 |
|  | In-M-cko | Ex-M-cko | .159 |
|  |  | Ctrl | .002 |
|  | Ctrl | Ex-M-cko | .221 |
|  |  | In-M-cko | .002 |

**Table2. The simple effects test result of the contextual fear memory among Ex-M-cko, In-M-cko and control.**
| group |  | Mean | Std. Error |
| --- | --- | --- | --- |
| Ex-M-cko | 1min | 49.541 | 5.028 |
|  | 2min | 57.568 | 5.605 |
|  | 3min | 48.568 | 6.716 |
|  | 4min | 46.277 | 6.946 |
|  | 5min | 46.741 | 5.320 |
| In-M-cko | 1min | 25.450 | 5.028 |
|  | 2min | 36.064 | 5.605 |
|  | 3min | 26.973 | 6.716 |
|  | 4min | 38.264 | 6.946 |
|  | 5min | 31.586 | 5.320 |
| Ctrl | 1min | 59.086 | 5.028 |
|  | 2min | 68.936 | 5.605 |
|  | 3min | 67.573 | 6.716 |
|  | 4min | 72.600 | 6.946 |
|  | 5min | 60.686 | 5.320 |

We did not see spontaneous behavioral seizures in the cko mutant mice. To test if the susceptibility to KA-induced seizures was changed, we injected KA (20mg/kg) intraperitoneally to observe acute behavioral seizure responses. The seizure score was evaluated every 20 minutes based on a modified Racine scale (29) totally for 120 minutes post injection (n=8/group). A Mixed Factorial ANOVA was performed to compare the seizure scores of female Ex-M-cko, In-M-cko and *Piga* ^WT^,Cre+ controls (Fig4.D). We observed a significant main effect for genotype: F(2, 30)=29.20, P<0.0001. Bonferroni corrected post hoc test showed that the seizure score of In-M-cko mutants was significantly higher than that of control, P<0.0001; and Ex-M-cko also demonstrated more seriously seizures than control, P=0.001. The multiple comparation of mean values (corrected by Bonferroni test) at each time point among the three genotypes is shown in Table 3A and B. A Mixed Factorial ANOVA was also performed to compare seizure score of male Th-H-cko mutants and “*Piga* ^flox/Y^, Cre-” control (Fig4.E). There was no significant difference between the two groups, F(1,14)=2.88, P=0.11. Therefore, deficit of *Piga* in either inhibitory or excitatory neurons increased the vulnerability of cko mutants to KA induced behavioral seizures.

**Table3.** The simple effects test result of the seizure score among Ex-M-cko, In-M-cko and control. (A)The Bonferroni test corrected p value of each comparison between the seizure score of two genotype groups at each time point. (B) The mean seizure score of each genotype at each time point.

| Time point(min) |  |  | Sig.<sup>b</sup> |
| --- | --- | --- | --- |
| Time-0 | Ex-M-cko | In-M-cko |  |
|  |  | Ctrl |  |
|  | In-M-cko | Ex-M-cko |  |
|  |  | Ctrl |  |
|  | Ctrl | Ex-M-cko |  |
|  |  | In-M-cko |  |
| Time-20 | Ex-M-cko | In-M-cko | .452 |
|  |  | Ctrl | 1.000 |
|  | In-M-cko | Ex-M-cko | .452 |
|  |  | Ctrl | .147 |
|  | Ctrl | Ex-M-cko | 1.000 |
|  |  | In-M-cko | .147 |
| Time-40 | Ex-M-cko | In-M-cko | .045 |
|  |  | Ctrl | .600 |
|  | In-M-cko | Ex-M-cko | .045 |
|  |  | Ctrl | .002 |
|  | Ctrl | Ex-M-cko | .600 |
|  |  | In-M-cko | .002 |
| Time-60 | Ex-M-cko | In-M-cko | .205 |
|  |  | Ctrl | .017 |
|  | In-M-cko | Ex-M-cko | .205 |
|  |  | Ctrl | .000 |
|  | Ctrl | Ex-M-cko | .017 |
|  |  | In-M-cko | .000 |
| Time-80 | Ex-M-cko | In-M-cko | .120 |
|  |  | Ctrl | .003 |
|  | In-M-cko | Ex-M-cko | .120 |
|  |  | Ctrl | .000 |
|  | Ctrl | Ex-M-cko | .003 |
|  |  | In-M-cko | .000 |
| Time-100 | Ex-M-cko | In-M-cko | .041 |
|  |  | Ctrl | .000 |
|  | In-M-cko | Ex-M-cko | .041 |
|  |  | Ctrl | .000 |
|  | Ctrl | Ex-M-cko | .000 |
|  |  | In-M-cko | .000 |
| Time-120 | Ex-M-cko | In-M-cko | .035 |
|  |  | Ctrl | .059 |
|  | In-M-cko | Ex-M-cko | .035 |
|  |  | Ctrl | .000 |
|  | Ctrl | Ex-M-cko | .059 |
|  |  | In-M-cko | .000 |
| Based on estimated marginal means |  |  |  |
| *. The mean difference is significant at the .05 level |  |  |  |
| b. Adjustment for multiple comparisons: Bonferroni |  |  |  |

| group |  | Mean | Std. Error |
| --- | --- | --- | --- |
| Ex-M-cko | 1 | 0.000 | 0.000 |
|  | 2 | 1.375 | .296 |
|  | 3 | 2.875 | .200 |
|  | 4 | 3.750 | .230 |
|  | 5 | 4.000 | .323 |
|  | 6 | 4.125 | .263 |
|  | 7 | 3.500 | .385 |
| In-M-cko | 1 | 0.000 | 0.000 |
|  | 2 | 2.000 | .296 |
|  | 3 | 3.625 | .200 |
|  | 4 | 4.375 | .230 |
|  | 5 | 5.000 | .323 |
|  | 6 | 5.125 | .263 |
|  | 7 | 5.000 | .385 |
| Ctrl | 1 | 0.000 | 0.000 |
|  | 2 | 1.125 | .296 |
|  | 3 | 2.500 | .200 |
|  | 4 | 2.750 | .230 |
|  | 5 | 2.250 | .323 |
|  | 6 | 2.375 | .263 |
|  | 7 | 2.125 | .385 |

### The deficit of *Piga* in excitatory neurons caused a decreased excitatory synaptic density and synapse size in hippocampus CA1

The brain structure of all the mutants including Ex-M-cko (Fig5.B), In-M-cko (Fig5.C) and Th-H-cko (Fig5.D) is comparable with that of WT controls(male Piga+/Y, Cre-, littermates of Th-H-cko) (Fig5.A). Furthermore, there was no obvious difference of the morphology of cerebella (Fig5.E~H). We expected that the complete deletion of *Piga* might cause the abnormal barrel formation in layer IV of somatosensory cortex in Th-H-cko mutants, and examined it by Nissl staining and cytochrome oxidase (CO) staining. However, we observed that barrels were normally formed in both Th-H-cko mutants (Fig5. J,L) and “*Piga* ^flox/Y^, Cre-” control (Fig5. I,K), and there was not a noticeable difference between them.

Our ISH result revealed that *Piga* is expressed most strongly in hippocampus excitatory neurons (Fig1. B). Based on that and the fact that Ex-M-cko showed a deficit in long-term fear memory, we investigated possible changes in excitatory synapses in hippocampus CA1 stratum radiatum by immunostaining with the postsynaptic marker PSD95. The results from Ex-M-cko and control groups are shown in Fig 5N and Fig 5.M, respectively. An unpaired t-test was conducted to compare the punctate number between Ex-M-cko and control groups (Fig5.O, n=24/group). Ex-M-cko (M=488.33,SE=8.7) showed a significant lower number than that of control (M=517.42,SE=8.9); t(46)=2.33, p=0.024. Due to the large sample size, a Z-test was performed to compare the puncta size of two groups (Fig5.P) (n=11720 puncta for Ex-M-cko; n=12418 puncta for control group), Z(critical two tail)=1.96, P<0.0001. These results suggested that the deficit of PIGA in excitatory neurons resulted in a decreased density and size of the excitatory synapses in CA1 stratum radiatum. In future work, it will be necessary to thoroughly examine the possible changes of both excitatory and inhibitory synapses in multiple brain regions in our mutants.

**Figure5.**
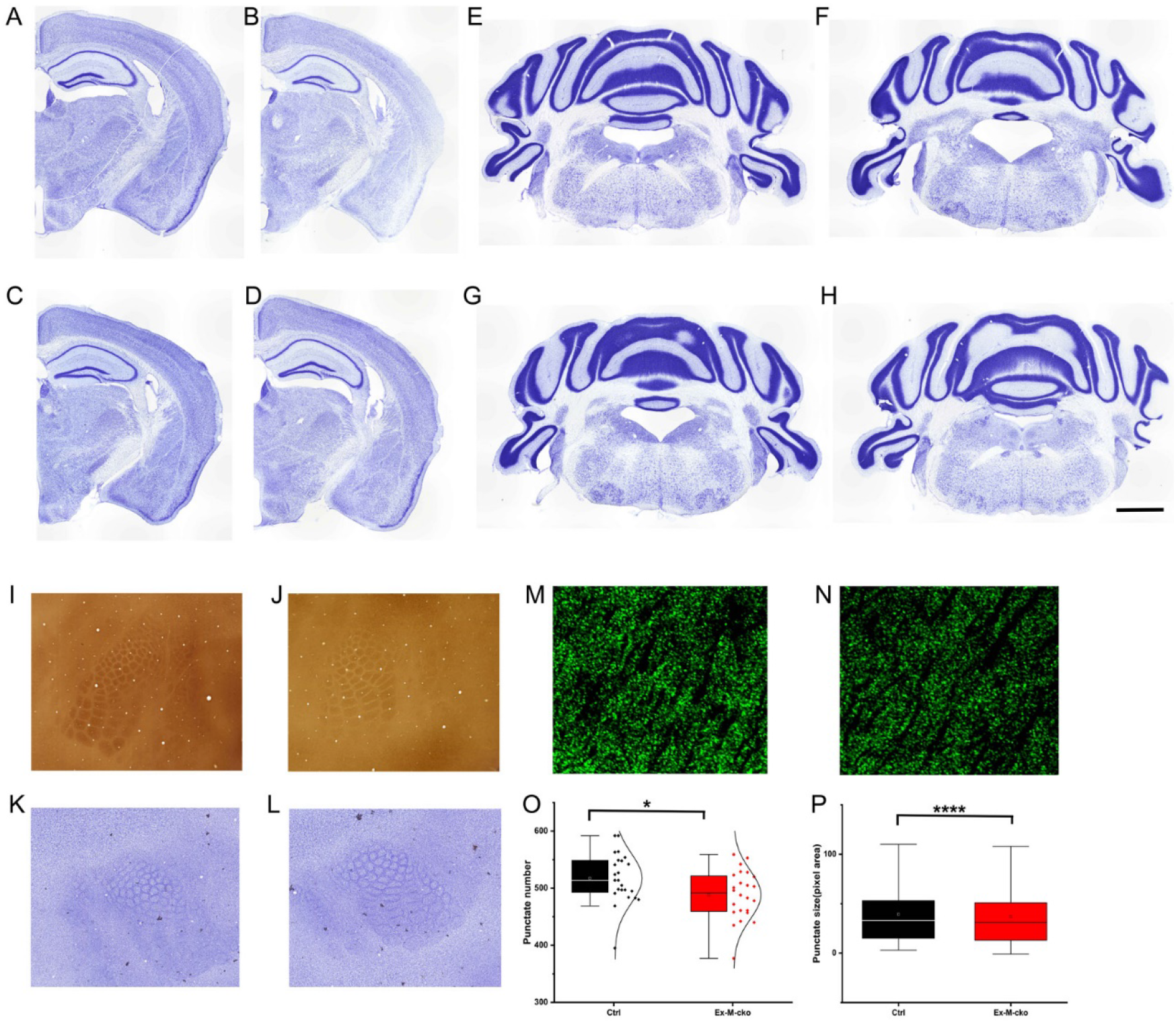
*Piga* mutants showed altered synaptic density and size in hippocampus CA1. (increase font for x axis) (A-H) Nissl staining of forebrain and cerebellum of WT(A,E), Ex-M-cko (B,F), In-M-cko (C,G) and Th-H-cko (D,H) revealed normal gross brain structure of mutants.(I-L) CO and Nissl staining of tangential sections of somatosensory cortex of *Piga*^flox^, Cre-male control (I,K) and Th-H-cko (J,L) suggested a normal barrel pattern in mutants. (M-P) PSD95 punctate number and punctate size was compared between Ex-M-cko mutants(M) and “*Piga*^WT^, Cre+” control mice(N). In the box charts, the boxes are determined by the 25th and 75th percentiles. The whiskers are determined by the 5th and 95th percentiles. Mean and medium values are shown as the small square and line inside the boxes. Each individual value is plotted beside the box (O,P). Scale bar: 1mm for (A-L); 12.5um for (M,N).

## Discussion

Since the loss function of PIGA completely abolishes GPI anchor production, *Piga* ^flox^ mouse line provided a powerful genetic tool to thoroughly explore the mechanisms underlying IGD encephalopathy in animal models. Using a *hCMV-Cre* mouse line, which expresses *Cre* before the preimplantation stage, Nozaki et al,. revealed that complete deletion of *Piga* totally perturbed organ formation in the early embryo stage of hemizygous cko mice (E9). Furthermore, partial knockout of *Piga* impaired development of neural tube and the face in the late embryo stage of mosaic cko from E12.5 to E17.5 (16). Lukas et al,. utilized a *Wnt1-Cre* mouse line to target *Piga* around E9.5 in neural crest cells and demonstrated that the deficit of *Piga* led to cleft lip/cleft and craniofacial hypoplasia, with hemizygous cko showing more severe phenotype than mosaic ones on E16.5 (20). In both cases the cko mice could not survive after birth. They also used the *Nestin-Cre* line to drive ablation of *Piga* in the central nervous system on E11.5, when the neural tube was already closed. None of the mutants survived past weaning, with 50% hemizygous cko not surviving 2 days after birth, and 50% of mosaic cko not surviving 19 days after birth. The brain size of mutants was smaller than that of WT (18). In the current study, we used *Emx1-Cre*, *Vgat-Cre* and *Pkcd-Cre* mouse lines to target ablation to telencephalon excitatory neurons, inhibitory neurons and thalamic neurons, respectively. We observed embryos of Ex-H-cko mutants with either microencephaly or normal appearance on E13.5, the reason of this discrepancy is unclear. Some In-H-cko embryos failed to develop any organs and appeared as an armorphous sphere on E13.5, while we also got In-H-cko embryos bearing similar appearance with the control embryos. Previous work reported that *Vgat* was expressed in a few sperm (30, 31), therefore the *Vgat-Cre* line may express *Cre* in a subpopulation of sperm. Neither the In-H-cko and Ex-H-cko mutants could survive after birth. All of the mosaic mutants and thalamus-specific hemizygous mutants (Th-H-cko) could survive as long as control mice, at least for 1 year. These data suggested that the short life span in Nestin-Cre induced *Piga* hemizygous mutants (Nestin-M-cko) (18) was more likely due to the deficits of GPI anchors in non-neuronal cells including astrocytes and oligodendrocytes.

There was a dramatic body weight loss in In-M-cko mice and a mild body weight loss in Ex-M-cko mutants. Previous work showed that Nestin-M-cko mutants were approximately half the weight of WT (18). Another study recently showed that a mouse model carrying *Pigv* hypomorphic mutation (*Pigv*^341E^) weighed about 60% of the WT control (32), which was similar to our In-M-cko mutants. Thus GPI-APs in inhibitory neurons may play an important role in regulating appetite and/or energy homeostasis. It has been reported that inhibitory neurons in the lateral hypothalamic area, ventral tegmental area and central nucleus of amygdala are involved in food intake behaviors (33). Future work examining the physiological changes of the inhibitory neurons in these brain regions of IN-M-cko mutants will help to identify the mechanisms underlying the reduced body weight. It is intriguing that limb-clasping was the most prominent motor phenotype in Nestin-M-cko (18), *Pigv*^341E^ (32) and our IN-M-cko mutants. Nestin-M-cko mutants also developed a severe tremor and ataxia, while *Pigv*^341E^ and IN-M-cko mutants did not. Limb-clasping could be related to dysfunction of the cerebellum and, indeed, Nestin-M-cko showed cerebellum atrophy and impaired Purkinje cell dendritic arborization (18). *Pigv*^341E^ showed normal-sized cerebellum and unchanged Purkinje cell dendrites. The cerebellum gross structure of our IN-M-cko mutants was unchanged as well. Limb-clasping is often seen in ataxia mouse models and is regarded as a marker of ataxia. However, consistent with our results, previous work also reported some mutants without overt ataxia showed very typical limb-clasping phenotype. These mutants include *Rai1* null mutants and *Ccnd1* null mutants with cerebellar atrophy, and *dt* mutants without cerebellum changes (34). The phenotypes of Nestin-M-cko (18), *Pigv*^341E^ (32) and our IN-M-cko mutants suggested that the mechanisms underlying limb-clasping, and ataxia and tremor might not overlap. Myelination in Nestin-M-cko was delayed during development (18) but not in *Pigv*^341E^ or our IN-M-cko mutants, further supporting that the impairment of myelination due to GPI deficit in oligodendrocytes was a possible reason causing serious tremor and ataxia in Nestin-M-cko. Future work is needed to more precisely elucidate the mechanism by assessing oligodendrocytes-specific cko mutants of *Piga*.

The most debilitating symptoms of IGD patients are epilepsy and intellectual disability. Our mutants had a normal life span, allowing us to investigate cognitive and seizure-related phenotypes. Fear conditioning has been widely used to test the ability of animal to learn and memorize an association between environmental/audial cues and an electrical shock. We tested our mutant mice with this paradigm and observed a significant decrease of fear response in the contextual memory test in IN-M-cko mutants and a decrease tendency in Ex-M-cko mutants. There was also a tendency that the cued memory of IN-M-cko was reduced. Contextual fear memory requires the coordination of hippocampus and amygdala. Although the gross structure of hippocampus and amygdala in the cko mutants was normal, a significantly smaller number and size of PSD95 puncta in hippocampus CA1 of Ex-M-cko mutant was detected. This suggests that the excitatory synapse number and size was altered in Ex-M-cko due to the deficit of GPI anchor synthesis. Spatial long-term memory impairment was reported in *Pigv*^341E^ hypomorphic mutant as well. In addition, there was a synaptic transmission defect in stratum radiatum of *Pigv*^341E^, including decreased EPSP amplitude, increased paired pulse facilitation and post-tetanic potentiation (32). Our ISH results revealed an enrichment of *Piga* in hippocampus excitatory neurons, and we previously observed facilitated PTP in stratum radiatum and impaired spatial memory in mutant mice deficient in synaptic adhesion GPI-AP, netrin-G2 (28, 35). All of these data implicate an important role of GPI-APs in the function of hippocampus pyramidal cells. KA animal models of epilepsy have been extensively used over the past decades because of their high level of similarity with human epilepsy. By systematic administration of KA, we observed an increased vulnerability to KA-induced behavioral seizures in both Ex-M-cko and IN-M-cko mutants. To our knowledge, two GPI-APs including netrin-G2 and contactin-2 have been reported to be related with epilepsy (36, 37). Specifically, contactin-2 null mice displayed spontaneous episodes of seizures (38) and netrin-G2/netrin-G1 double knockout mice were more susceptible to KA-induced seizures (unpublished data). It has been commonly assumed that an imbalance between the excitatory and inhibitory synaptic transmission in the brain initiates seizure activity. Future studies will be necessary to thoroughly examine the morphological, molecular and physiological changes of both excitatory and inhibitory neurons in different subregions of hippocampus and amygdala in our mutants.

It is noteworthy that although we obtained Th-H-cko mutants and achieved essentially complete deletion of *Piga* in thalamic neurons, we did not observe any obvious abnormality of these mice. Their barrel cortex formed normally. The *Pkcd-Cre* induced recombination postnatally and then it took a couple of weeks to reach the maximal level(unpublished data). This timing of recombination might explain the minor effects resulted from *Pkcd-Cre* induced *Piga* deletion.Netrin-G1 is a GPI-AP which is highly enriched in thalamus. Netrin-G1 knockout mice showed normal barrel cortex (27), and thalamic-specific cko of netrin-G1 performed similarly to control littermates in all the behavioral paradigms we tested ((23) and unpublished data). In addition, a previous study revealed that T cells deficient in GPI-APs were functionally competent to respond to T cell receptor stimulation both in vitro and in vivo (39). Therefore GPI-APs are dispensable for some functions in certain cell populations. Future research will be needed to examine more detailed and delicate features of the thalamus related functions of Th-H-cko mutants.

Different cell populations in the nervous system have different expression profiles of GPI-APs. To date, many GPI-APs have been reported to be expressed in neurons, including Thy-1, TAG-1, NCAM120, Contactin,T-cadherin, LAMP, OBCAM, Neurotrimin, Kilon, ephrin-A, semaphoring-7A, glypicans, netrin-Gs, NgRs, MDGAs, GFRα1, PrP^C^, IgLON, CPG15/Neuritin. These molecules perform diverse functions in axonal fasciculation, growth and guidance, neuronal migration, and synapse formation and plasticity (40–45). A few GPI-APs have been discovered to be expressed in glial cells. For examples, Glipican4 and 6 are expressed by astrocytes, with Glipican4 expression enriched in the hippocampus and Glipican6 in the cerebellum, and they play roles in synapse formation and synaptic plasticity (46). Contactin, TAG-1 and OMgp are expressed in oligodendrocytes and regulate neurite growth, myelination, nodal/paranodal domain organization and synaptic plasticity (40, 41, 43, 45). It is interesting that *Pigv*^341E^ mutants showed similar phenotypes to our IN-M-cko mutants but not Nestin-M-cko. In *Pigv*^341E^ mutants, all the cells including oligodendrocytes carry the mutation *Pigv*:1022C>A (p.A341E). However, there was no obvious white matter deficits observed in Nestin-M-cko. All the IGD patients reported so far carried mutations of a molecule involved in GPI anchor synthesis or maturation. The symptoms were variable and were resulted from the sum effect of GPI-APs deficits in affected cell populations. We hypothesize that the expression of a GPI synthesizing molecule was regulated by different elements in different cell types or even subtypes. The point mutation 1022C>A in *Pigv*^341E^ mouse may mainly affect the expression of *Pigv* in neurons but not in oligodendrocytes. It remains to be tested explicitly whether specific or common transcription factors regulate the expression of a certain GPI synthesizing molecule in different cell populations. Another intriguing point is that in human beings, all *PIGA*-IGD patients are male. Their mothers, who are heterozygous carriers of PIGA mutation, were phenotypically mosaic at the cell level but don’t show clear symptoms as previously described (3). In mice, the female heterozygous carriers of *Piga* mutations demonstrated clear deficits. It has been reported that the regulation of x-linked gene expression is quite different between human and mouse(47), future work will be needed to investigate the mechanisms underlying the species difference of phenotypes we observed here. Nontheless, Nestin-M-cko, *Pigv*^341E^, IN-M-cko, Ex-M-cko mutants are the only GPI-anchor deficient mouse models that can survive after birth and mirror the neurological symptoms of IGD patients. Together with the coming oligodendrocyte-specific *Piga* cko, they will be useful animal models to help elucidate the molecular mechanisms of IGD and test the novel therapeutic approaches for the intractable symptoms of IGD patients.

## Materials and Methods

### Animals

All of the mice were housed in a temperature and humidity controlled, pathogen free environment maintained on a 12:12h light/dark cycle, with food and water *ad libitum*. *Piga*^flox^ mice (17) (RBRC06211) were obtained from Dr. Junji Takeda, Dr. Taroh Kinoshita and RIKEN BRC. *Emx*-Cre, *Vgat*-Cre and *Pkcd*-Cre mice were described previously (21–23). The hemizygous and mosaic cko mice were obtained by breeding female *Piga*^flox/+^ mice with male *Cre* mice, and were housed in groups with mixed genotype. Unless specified, Ex-M-cko and IN-M-cko mutants aged between 16 weeks to 32 weekes were compared with their female littermates (*Piga*^+/+^,Cre+), Th-H-cko mutants aged between 16 weeks and 32 weekes were compared with their male littermates (*Piga*^flox/Y^,Cre-). All assays were performed by experimenters blind to the animals’ genotype.

### Genotyping and PCR analysis of neuronal specific disruption of Piga

For genotyping by using tail DNA, the PCR primer pairs were as follows: Cre1: 5’-TCGACCAGGTTCGTTCACTC-3’and Cre2: 5’-TGACCCGGCAAAACAGGTA-3’ amplified bands of 304 bp for Cre recombinase. Primer1: 5’-ACCTCCAAAGACTGAGCTGTTG-3’ and Primer2: 5’-CCTGCCTTAGTCTTCCCAGTAC-3’ amplified bands of 420bp for *Piga* floxed allele, and 250bp for wild type allele.

For assessing the deletion efficiency of *Piga* in neuronal populations, we followed a protocol as previously described (24) with some modifications. Briefly, hippocampus, stratum, thalamus and cortex were dissected from 8 brains/genotype. Single cell suspension was obtained by trypsin digestion and cell strainer filtering and then briefly fixed in 50% ethanol. The cells were stained by 1^st^ antibody anti-NeuN (Millipore) and 2^nd^ antibody Alexa Fluor 488 donkey anti-mouse IgG (Molecular Probes) and then sorted by BD FACSAria II. And the three PCR primers were as follows: Primer1: 5’-ACCTCCAAAGACTGAGCTGTTG-3’; Primer2: 5’-CCTGCCTTAGTCTTCCCAGTAC-3’ and Primer3: 5’-TGTGGGTTTCAGTTCATTTCAGA-3’ amplified bands of 420bp for floxed allele, 250bp for wild type allele, and 550bp for disrupted configuration.

### *In situ* hybridization

The probe sequence was 790 bp from the non-coding region of *Piga* exon 6. It was obtained by PCR from mouse hippocampus cDNA library, the following primers were used: Forward: GGGATAATGGTTTAGCCACTCA and Reverse: GAAGAGCAGCTGGTTTTGAGAT. Preparation of digoxigenin-labeled antisense riboprobes, and hybridization of free-floating sections were performed according to previously described procedures(23).

#### Immunohistochemistry

The netrin-G1 and PSD95 antibodies (State sources), and the immunohistochemistry procedure on free-floating sections were described in previous work (27, 35).

### Histology

The procedure of Nissl staining and CO staining were described previously (21).

### Limb-clasping scoring

Mice were held by the tail for 5 to 10 seconds. During tail suspension, limb clasping behavior was manually evaluated with following standards: 0-No limb clasping, Normal escape extension. 1-Either hind limbs or forelimbs incomplete splay and loss of mobility. 2-Either hind limbs or forelimbs clasping together and loss of mobility. 3-Single side hind limb and forelimb clasping together and loss of mobility. 4-Three limbs clasping together and loss of mobility. 5-Four limbs clasping together and loss of mobility. 6-Four limbs clasping together, body forms a ball shape and loss of mobility.

### Fear conditioning test

Contextual and cued fear conditioning test were performed as previously described (23).

### Behavioral seizure scoring

KA-induced behavioral seizures were scored as previously described (48).

### Punctate number and size quantification

Four mice from each group were analyzed. 3 continuous sections around the level of Bregma −2mm were obtained from each mouse, totally 24 images from the similar positions of stratum radiatum of each genotype were caught by Olympus FV3000 under the same condition. The punctate number and punctate size from each frame(2700um^2^) were measured with Metamorph Software.

### Statistics

All analyses were conducted using SPSS Statistics21. For body weight comparation among three female groups, ANOVA was used, following by Bonferroni corrected post hoc testing. For body weight comparation between two male groups, unpaired t-test was used. For limb-clasping score comparation among three groups, Kruskal-Wallis test was used followed by Bonferroni corrected post hoc testing. For limb-clasping score comparation between two male groups, Man-Whitney U test was used. For all time series data of behavioral analysis, Mixed Factorial ANOVA was used to compare among groups, followed by post hoc testing via the Bonferroni correction. In addition, simple effects test at each time point via the Bonferroni correction was applied when group effects existed and the data were summarized in Table2 and Table3. For the comparation between two groups with sample size bigger than 30, two tailed Z-Test was applied.

## Acknowledgements

This project was supported by the RIKEN Incentive Research Project (100226201701100443), the Brain Science Project, Center for novel science initiatives, National institutes of natural sciences (BS291003), the RIKEN Aging Project (10026-201701100263-340120), the JSPS Kakenhi Grant-in-Aid for Young Scientists (17K18362), the JSPS Kakenhi Grant-in-Aid for Challenging Research (19K21807), and the International Education and Research Laboratory Program of University of Tsukuba. We thank Dr. Taroh Kinoshita for very informative and inspiring discussion.

## Contributions

Q.Z. conceived and designed the research; L.K., M.T., V.B., S.T., Y.N. and Q.Z. conducted the experiments and analyzed the data. M.K. and N.M. provided all the necessary information of clinical patients carrying *PIGA* mutations. J.K. provided *Piga*^flox^ mouse. S.I., S.O., and L.Y. provided the experimental facility and resources. Q.Z., N.M., and L.Y. interpreted data. Q.Z. drafted the manuscript. N.M., S.I and L.Y. commented on and edited the manuscript.

